# Gestational arsenite exposure alters maternal postpartum heart size and induces Ca^2+^ handling dysregulation in cardiomyocytes

**DOI:** 10.1101/2024.09.25.615085

**Authors:** Nicole Taube, Morgan Steiner, Obialunanma V. Ebenebe-Kasonde, Raihan Kabir, Haley Garbus-Grant, Sarah-Marie Alam El Din, Emily Illingworth, Nadan Wang, Brian L. Lin, Mark J. Kohr

## Abstract

Cardiovascular disease is the leading cause of mortality in the US. Studies suggest a role for environmental exposures in the etiology of cardiovascular disease, including exposure to arsenic through drinking water. Arsenic exposure during pregnancy has been shown to have effects on offspring, but few studies have examined impacts on maternal cardiovascular health. While our prior work documented the detrimental effect of arsenic on the maternal heart during pregnancy, our current study examines the effect of gestational arsenic exposure on the maternal heart postpartum. Timed-pregnant wild-type (C57BL/6J) mice were exposed to 0, 100 or 1000 µg/L sodium arsenite (NaAsO2) via drinking water from embryonic day 2.5 (E2.5) until parturition. Postpartum heart structure and function was assessed via transthoracic echocardiography and gravimetric measurement. Hypertrophic markers were probed via qRT-PCR and western blot. Isolated cardiomyocyte Ca^2+^-handling and contraction were also assessed, and expression of proteins associated with Ca^2+^ handling and contraction. Interestingly, we found that exposure to either 100 or 1000 µg/L sodium arsenite increased postpartum heart size at P12 vs. non-exposed postpartum controls. At the cellular level, we found altered cardiomyocyte Ca^2+^-handling and contraction. We also found altered expression of key contractile proteins, including α-Actin and cardiac myosin binding protein C (cMyBP-c). Together, these findings suggest that gestational arsenic exposure impacts the postpartum maternal heart, possibly inducing long-term cardiovascular changes. Furthermore, these findings highlight the importance of reducing arsenic exposure during pregnancy, and the need for more research on the impact of arsenic and other environmental exposures on maternal heart health and adverse pregnancy events.

**New & Noteworthy:** Gestational exposure to sodium arsenite at environmentally relevant doses (100 and 1000 µg/L) increases postpartum heart size, and induces dysregulated Ca^2+^ homeostasis and impaired shortening in isolated cardiomyocytes. This is the first study to demonstrate that gestational arsenic exposure impacts postpartum heart structure and function beyond the exposure period.

## INTRODUCTION

Exposure to inorganic arsenic (iAs) is a widespread public health concern, with an estimated 140 million people worldwide exposed to levels of iAs in drinking water in excess of the World Health Organization’s recommended limit of 10 µg/L^1,2^. iAs occurs naturally in the earth’s crust, but anthropogenic processes such as mineral extraction, waste processing, pesticide application and additives to poultry and swine feed have mobilized iAs into drinking water supplies^3,4^. Countries such as China, Hungary, and the United States have all found high levels of iAs in groundwater, primarily affecting regions dependent on well water^5^. Of the two forms of iAs, trivalent arsenic has been found to be the most toxic, with pentavalent arsenic being excreted or transformed into trivalent arsenic^6,7^. iAs has been linked to the etiology of many disease states, including cancer, chronic respiratory disease, and cardiovascular disease (CVD)^8–16^. Specifically, trivalent iAs exposure is associated with hypertension, atherosclerosis, coronary heart disease, stroke, and diabetes^5,13,14,17–20^. Our group previously reported that chronic exposure to iAs induced sex-specific pathological remodeling of the adult heart and altered susceptibility to ischemic injury,^21,22^ but few studies have examined how iAs impacts the cardiovascular system under physiologic stress, including the maternal heart after pregnancy.

In healthy pregnancies, cardiovascular changes are normally well tolerated, but in some cases adverse pregnancy events and cardiovascular complications may arise. In a healthy pregnancy, increased plasma volume drives reversible cardiac hypertrophy^23^, leading to an increase in wall thickness and left ventricular wall mass by approximately 52% compared to pre-pregnancy values^24^. During the postpartum period, the heart remodels back to the pre-pregnant size in 7 to 14 days in a mouse^25,26^ or up to a year in humans^24,27^. Additionally, cardiovascular complications during pregnancy may also lead to postpartum CVD^28^. Cardiovascular complications are the leading cause of maternal mortality and can result from preeclampsia or peripartum cardiomyopathy^28–30^. While these complications can result from comorbidities established prior to pregnancy, such as diabetes or hypertension, complications arise in many pregnancies which are seemingly idiopathic^29^. As such, the environment is thought to contribute to these adverse events during pregnancy. For example, exposure to air pollution, heavy metals and perfluorinated chemicals have all been correlated with preeclampsia^31–34^. However, more research is needed to understand how environmental factors are involved in adverse cardiovascular events during the perinatal period and to elucidate underlying mechanisms.

Our group previously reported that gestational exposure to iAs abrogated normal maternal cardiac growth during late pregnancy^35^. Here, we used the same model of gestational iAs exposure to determine potential impacts to postpartum cardiac structure and function at postnatal day 5 (P5) and isolated cardiomyocyte Ca^2+^ handling and contraction at P14/15. We report that exposure to iAs during gestation alone results in increased heart size postpartum, as well as dysregulated Ca^2+^ handling and contraction at the level of the isolated cardiomyocyte.

## METHODS

### Animals and Arsenic Exposure

This investigation conforms to the Guide for the Care and Use of Laboratory Animals published by the National Institutes of Health (NIH; Publication No. 85–23, Revised 2011) and was approved by the Institutional Animal Care and Use Committee of Johns Hopkins University. Timed pregnant female C57BL/6 J mice were purchased from Jackson Laboratory (Bar Harbor, ME). Mice (2 per cage) were housed under specific pathogen-free conditions and maintained on AIN-93G chow (Research Diets, New Brunswick, NJ) and Pure Life water (Nestle Waters North America, Stamford, CT). Research Diets has reported levels of iAs in AIN-93G chow to be below the limit of detection, and our lab has previously confirmed levels of iAs in Pure Life water to be below the limit of detection using inductively coupled plasma mass spectrometry measurements^35–37^. Pregnant and non-pregnant female mice (6-12 weeks of age) were given Pure Life water containing 0 (control), 100 or 1000 µg/L sodium arsenite (NaAsO_2_) beginning at embryonic day (E) 2.5 and continuing until parturition or for 16 days (E18.5) for non-pregnant mice^35^. Following the guidelines provided by the Food and Drug Administration (FDA), allometric scaling was be used to determine human to rodent equivalency dose and we found that our 100 and 1000 µg/L sodium arsenite doses are equivalent to 8.3 µg/L and 83.33 µg/L, respectively^38^. Fresh water bottles containing control or NaAsO_2_ drinking water were replenished every 2–3 days to maintain concentrations of iAs and minimize oxidation. To reduce stress during pregnancy, dams were only handled when necessary (i.e., weight measurement, cage change, etc.). Furthermore, dams which cannibalized their young and therefore not lactating postpartum, were excluded from this study. Body weight, and food and water consumption did not significantly differ between treatment groups throughout the exposure as reported in our prior study^35^.

### Tissue Harvest and Collection

Mice were anesthetized with a mixture of ketamine (90 mg/kg; Hofspira) and xylazine (10 mg/kg; Sigma-Aldrich) via intraperitoneal injection and anticoagulated with heparin (Fresenvis Kabi, Lake Zurich, IL) prior to tissue collection. Hearts were excised, cannulated on a Langendorff apparatus, and perfused in a retrograde manner with Krebs-Henseleit buffer (95% O2, 5% CO2; pH 7.4) under constant pressure (100 cmH_2_O) and temperature (37 °C) as previously described for 5 min to washout blood^35,37^. Buffer consisted of (in mmol/L): NaCl (120), KCl (4.7), KH_2_PO_4_ (1.2), NaHCO_3_ (25), MgSO_4_ (1.2), d-Glucose (14), and CaCl_2_ (1.75). Following perfusion, hearts were weighed and either sectioned for histology, or equally halved, and snap-frozen in liquid nitrogen and stored at −80 °C for molecular studies. Heart size was also normalized to tibia length to account for any age-dependent differences. Tissue was collected at P12. Cross-fostering of offspring was not performed in this study, and dams were kept with their respective litters until P12.

### Transthoracic Echocardiography

All echocardiography procedures were performed by an investigator blinded to experimental conditions using a preclinical ultrasound imaging system (Vevo 2100; FUJIFILM VisualSonics, Toronto, ON, Canada) with a 40-MHz linear transducer, as previously described^35,37^. Echocardiography was performed on mice under anesthesia. Mice were induced with 3-4% isoflurane and maintained at 1-2% isoflourane thereafter. The M-mode echocardiogram was acquired from the short-axis view of the left ventricle at the level of the midpapillary muscles (200 m/s sweep speed). Semiautomated continuous tracing of the left ventricular walls from this axis view was used to measure, calculate, or extrapolate the following cardiac parameters using 3-5 cardiac cycles: left ventricular posterior wall thickness at end diastole (LVPWd), left ventricular posterior wall thickness at end systole (LVPWs), left ventricular internal diameter at end diastole (LVIDd), left ventricular internal diameter at end systole (LVIDs), percent fractional shortening (FS), percent ejection fraction (EF), heart rate (H), and cardiac output (CO). Parameters that were calculated were done so using the LVTool function using the Vevo LAB software (VisualSonics, Fujifilm). Echocardiography was performed at P5.

### RNA Isolation, Extraction, and cDNA Conversion

Hearts were homogenized with TRIzol (1 mL, Ambion) using a bead-mill tissue homogenizer (2 × 30 s cycles, 0 °C, 7200 rpm; Precellys Evolution 24). Lysates were mixed with chloroform (200 μL, Thermo Fisher), incubated (5 min, 25 °C), and centrifuged (12,000 g, 15 min, 4 °C) for phase separation. RNA collected from the upper phase was precipitated by incubation (10 min, 25 °C) with isopropyl alcohol (500 μL) and centrifugation (12,000 g, 8 min, 4 °C). RNA pellets were washed with 75% ethanol, centrifuged (12,000 g, 10 min, 4 °C), and air-dried under a laminar flow hood (30 min). Following RNA solubilization (100 μL, DEPC-treated water), RNA concentration and purity (A260/A280 range 1.98 to 2.06) were measured via spectrophotometry (1 µL sample, NanoDrop 100, Thermo Fisher). RNA was subsequently converted to cDNA per the manufacturer’s instructions (High-Capacity cDNA Reverse Transcription Kit, 4,368,814, Thermo Fisher). Briefly, the reverse transcription reaction mix was prepared on ice, RNA (2 μg) was added, and the samples were run on a thermocycler (60 min, 37 °C; 5 min, 95 °C; held, 4 °C; Applied Biosystems); cDNA was stored at −20 °C until use.

### Quantitative RT-PCR

Expression levels of mRNA transcripts were measured using generated cDNA, a PCR master mix (TaqMan Fast Advanced Master Mix, Applied Biosystems) and the following validated primers (TaqMan, Applied Biosystems) on a thermocycler (2 min, 50°C; 2 min, 95°C; (1 s, 95°C; 20 s, 60°C) × 40), (QuantStudio 3, Applied Biosystems, Thermo Fisher) in a 96-well plate (USA Scientific, Beltsville MD): *Myh6* (Mm00440359_m1), *Myh7* (Mm00600555_m1), *Cacna1d* (Mm01209927_g1), *RyR2* (Mm00465877_m1), *Atp2a2* (Mm01201431_m1), and *Gapdh* (Mm99999915_g1). Expression was determined using the ΔΔCT method and normalized to *Gapdh*, which did not change in cycle time with iAs treatment.

### Heart Homogenization and Protein Isolation

Hearts were homogenized in cell lysis buffer (1 mL; Cell Signaling Technology, Danvers, MA) supplemented with a protease and phosphatase-inhibitor cocktail (Cell Signaling Technology) using a hard tissue lysing kit (Precellys CK28 Lysing Kit, Bertin Instruments) with a bead-mill tissue homogenizer (2 x 30 s cycles, 0 °C, 72000 rpm; Precellys Evolution 24, Bertin Instruments). Supernatant was collected as total crude homogenate; protein concentration was determined via Bradford assay and aliquots of homogenate were stored at −80 °C.

### Western Blot

Protein homogenates (30 µg) were separated (20 min, 75 V; 100 min, 175 V) on a gradient Bis-Tris SDS-PAGE gel (4-12%, NuPAGE; Thermo Fisher, Carlsbad, CA) and transferred (90 min, 220 mA) to a nitrocellulose membrane (Thermo Fisher). Every gel included two molecular-weight markers for separate regions of interest (High Range Color-Coded Prestained Protein Marker; Cell Signaling Technology; and Novex Prestained Protein Standard, Thermo Fisher) on opposite ends of the gel. Total protein served as the loading control by covalently labeling lysine residues (No-Stain Protein Labeling Reagent, Thermo Fisher) following the manufacturer’s instruction and visualized by fluorescence imaging (488 nm). Membranes were blocked (1h) with bovine serum albumin (5% wt/vol, Sigma-Aldrich) in Tris-buffered saline with Tween-20 (0.1%), and subsequently incubated (overnight, 4 °C) with primary antibodies for p-Akt (S473) (1:1000, Rabbit mAb, Cell Signaling, 4060S), Akt (pan) (1:1000, Rabbit mAb, Cell Signaling, 4691S), p-eNOS (S1177) (1:1000, Rabbit mAb, Cell Signaling, 9571S), eNOS (1:500, Rabbit mAb, Santa Cruz, sc-376751), p44/42 MAPK (ERK1/2) (1:1000, Rabbit mAb, Cell Signaling, 9102S), p-p44/42 MAPK (T202/Y204) (1:1000, Rabbit Ab, Cell Signaling, 9101S), SERCA2a (1:500, Goat Ab, Santa Cruz, sc-8094), p-PLBN (S16/Thr17) (1:1000, Rabbit Ab, Cell Signaling, 8496S), total PLBN (1:1000, Rabbit Ab, Cell Signaling, 14562S), LTCC α-2 subunit (1:1000, Rabbit Ab, Abcam, ab253190), Mybp-c (1:1000, Rabbit Ab, Santa Cruz Technologies, sc-137180), and α-Actin (1:1000, Mouse Ab, Proteintech, 23660-1-AP). After primary incubation, membranes were incubated with secondary antibodies (Anti-rabbit, - mouse or -goat IgG, HRP-linked Ab, Thermo Fisher) and visualized using chemiluminescence substrate (SuperSignal West Pico PLUS, Thermo Fisher) and an iBright imager (Thermo Fisher). Membranes were stripped with Reblot Plus Mild Solution (Millipore) when necessary to visualize total protein when examining phosphorylation status.

### Cardiomyocyte Ca^2+^ transients and sarcomere shortening

Immediately after cannulation, hearts were perfused, and cardiomyocytes were isolated as previously described^39^. Briefly, hearts were perfused with buffer containing Type II Collagenase (Worthington) and protease (Sigma) for approximately 9 mins. Following digestion, hearts were minced and triturated gently. The cell suspension was then filtered and centrifuged at 800 RPM for 1 minute. Isolated cardiomyocytes were then rinsed and stored in Tyrode buffer (pH 7.4) containing the following (in mmol/L): NaCl (140), KCl (5), MgCl_2_ (1), HEPES (10), and glucose (5.5), and were gradually stepped up every 10 minutes to a final [Ca^2+^] of 1 mmol/L. Viability of cardiomyocytes was estimated to be 70-90%, and only live cardiomyocytes were used for measurements. Ca^2+^ transients and sarcomere shortening were then measured using an inverted microscope (Nikon Eclipse TE-2000U) and a custom-built IonOptix system with IonWizard software as previously described^40^. Cardiomyocytes were stimulated at 1 Hz and maintained at 37°C in Tyrode buffer for all measurements. Fura-2-AM was excited at 340 and 380 nm alternating at 250 Hz and emission recorded at 510 nm by a single photon multiplier tube. Background was subtracted from Fura-2 readings and the results filtered with a Lowpass Butterworth Filter (cutoff frequency of 10 Hz, 2 poles). Multiple Ca^2+^ signal transients (10-15) per cell were averaged for analysis. The fluorescence ratio value is shown as F/F_0_ (fluorescence normalized to baseline 340/360), and the fit used to calculate shortening tau is standard from the Ionoptix Monotonic Transient Analysis^41^. Cardiomyocytes were used for functional measurements within 6 hours of isolation. For this study, 40-80 cardiomyocytes were used per heart, with four hearts per exposure group. For statistical analysis, a two-way ANOVA was performed using n=40-80 cardiomyocytes per heart.

### Statistical Analysis

Sample sizes of mice for each experiment were estimated a priori via power analysis (power = 0.80, effect size = 0.25, alpha = 0.05) based on data generated in previous studies from our group. Mice were randomized during collection to minimize batch effects. Data was analyzed using GraphPad Prism (La Jolla, CA). Statistical outliers were identified using the ROUT method (Q=1%). Statistical comparisons between groups were determined using an ordinary one-way ANOVA with Dunnett’s or Sidak’s multiple comparisons test, a two-way ANOVA with Tukey’s multiple comparisons test or a two-tailed Mann-Whitney test as appropriate. Significance was set at p < 0.05.

## RESULTS

### iAs exposure increases maternal heart size postpartum, but does not alter function

Our previous report demonstrated that prenatal exposure to iAs blunted maternal heart growth during late pregnancy^35^. In the current study, we investigated if these effects persisted postpartum. Using gravimetric measurement, we found that exposure to both 100 µg/L and 1000 µg/L iAs increased postpartum heart size at P12 compared to controls (Fig. 1A). Transthoracic echocardiography was also performed on dams at P4 and P5, but we found that exposure to 100 µg/L and 1000 µg/L iAs did not alter ejection fraction (Fig. 1B) or fractional shortening (Fig. 1C) compared to non-exposed postpartum controls. We also observed no change in cardiac output (Fig. 1D), stroke volume (Fig. 1E), heart rate (Fig. 1F), end diastolic volume (1G), end systolic volume (1H) or left ventricular internal diameter during diastole (LVID;d) (Fig. 1I).

**Figure 1:**
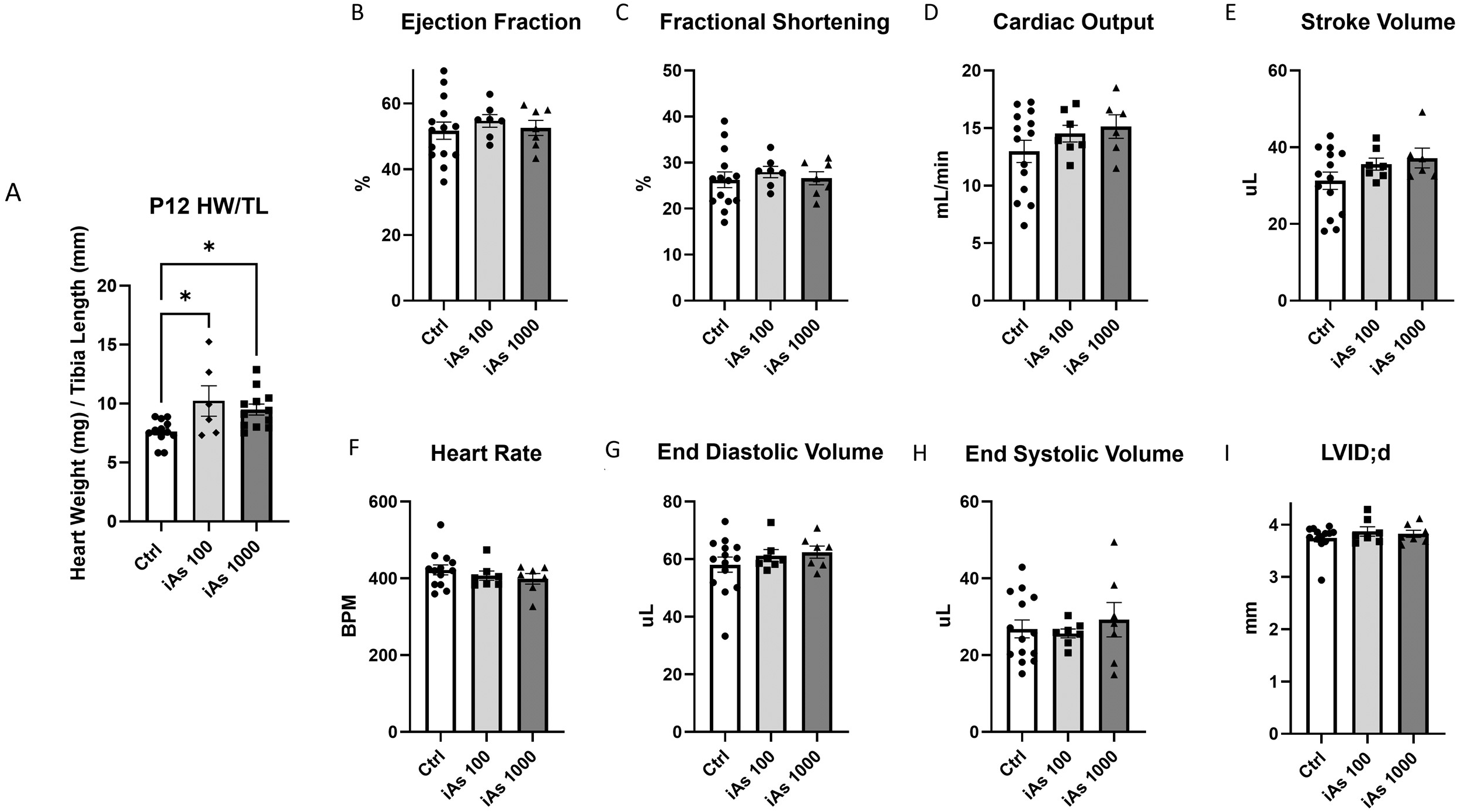
Gestational iAs exposure increases maternal heart size postpartum but does not impact function. (A) Heart weight divided by tibia length in postpartum females following gestational exposure to 100 µg/L or 1000 µg/L iAs exposure compared to non-exposed controls (n=6-12 mice/group; *= P < 0.05 vs. control). Significance was determined by one-way ANOVA with Tukey’s multiple comparisons test. (B-I) Cardiac parameters as measured by transthoracic echocardiography including: ejection fraction (B), fractional shortening (C), cardiac output (D), stroke volume (E), heart rate (F), end diastolic volume (G), end systolic volume (H), left ventricular internal diameter in diastole (LVID;d;) in iAs-exposed or non-exposed postpartum females (n=7-14 mice/group). Significance was determined by one-way ANOVA with Tukey’s multiple comparisons test.

### iAs exposure alters cardiac hypertrophic signaling markers postpartum

Since we observed an increase in heart size, we next probed major markers of physiologic and pathologic hypertrophy in dams exposed to 100 µg/L iAs and controls. Furthermore, all subsequent mechanistic experiments were performed at the 100 µg/L iAs dose because this was the lowest dose where we observed a change in heart size. We found that total endothelial nitric oxide synthase (eNOS) expression was unchanged with iAs exposure in postpartum hearts compared to non-exposed postpartum controls (Fig. 2A), but phosphorylation of the active site (Ser1177) was significantly decreased in postpartum hearts compared to non-exposed postpartum controls (Fig. 2B). Additionally, expression of protein kinase B (Akt) was downregulated with iAs exposure in postpartum hearts compared to non-exposed postpartum controls (Fig. 2C), but phosphorylation at the active site (Ser437) was not altered (Fig. 2D). We also examined extracellular signal-regulated kinase 1/2 (ERK-1/2) but found that iAs exposure did not alter the expression of ERK-1/2 (Fig. 2E) or phosphorylation at the active site (Thr202/Tyr204) (Fig. 2F). Additionally, we probed transcription levels of α-myosin heavy chain (*Myh6*) and β-myosin heavy chain (*Myh7*) for evidence of pathological remodeling but found that iAs did not alter transcription of either target (Figs. 2G, 2H).

**Figure 2:**
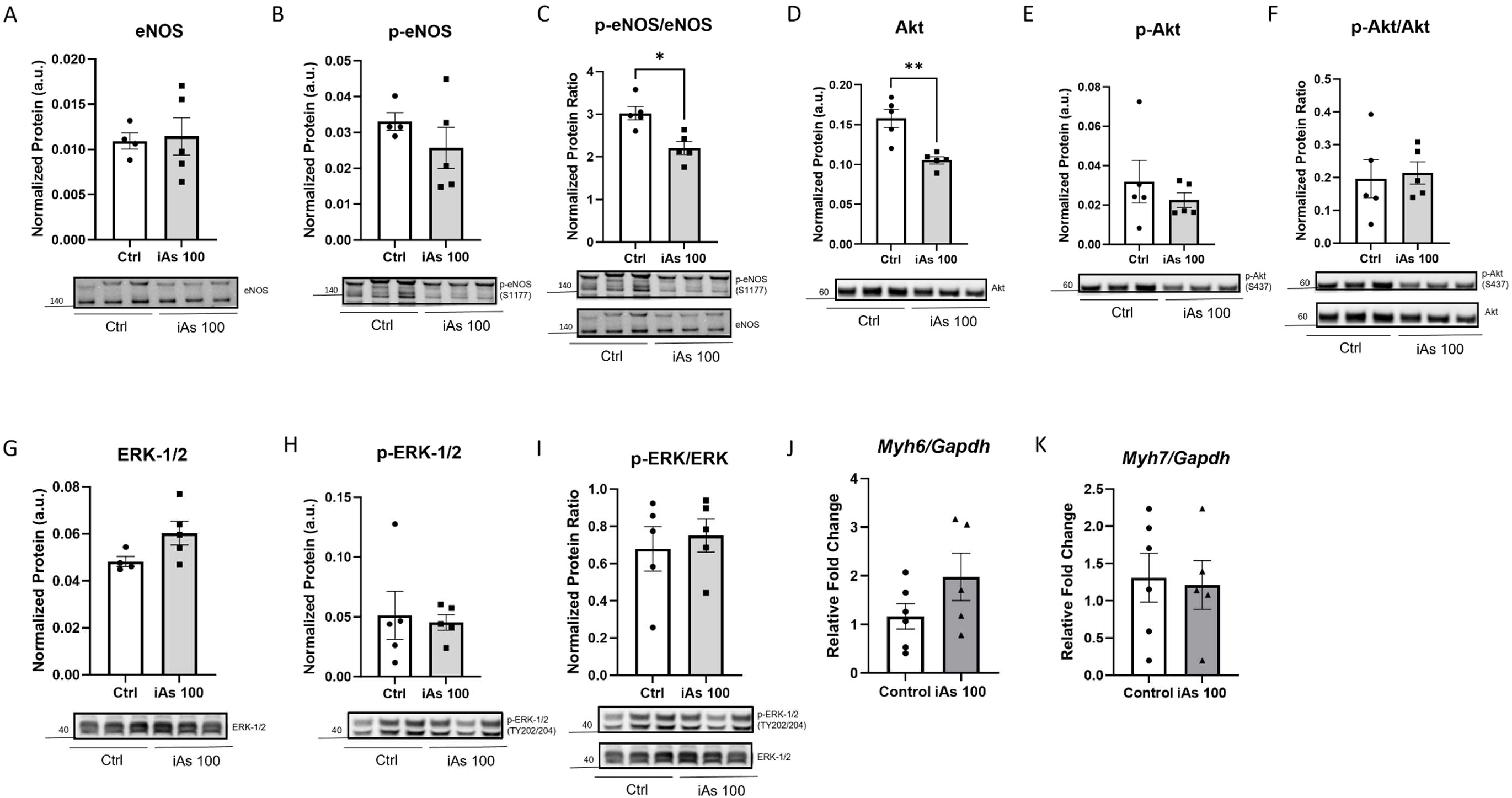
Gestational iAs exposure alters expression and phosphorylation of hypertrophic signaling markers at P12. Myocardial protein expression of endothelial nitric oxide synthase (eNOS) (A), phosphorylated eNOS at Ser1177 (B), p-eNOS relative to total at Ser1177 (C), protein kinase B (Akt) (D), phosphorylated Akt at Ser437 (E), p-Akt relative to total at Ser437 (F), extracellular signal-regulated kinase 1/2 (ERK-1/2) (G), p-ERK-1/2 at Thr202 and Tyr204 (H), p-ERK1/2 relative to total (I), and myocardial transcription levels of α-myosin heavy chain (*Myh6*) (J) and β-myosin heavy chain (*Myh7*) (K) in iAs-exposed or non-exposed postpartum females. (*n*=4-5 hearts/group or 8-10 hearts/group; *p<0.05, **p<0.01 vs. control). Significance was determined by Mann-Whitney.

### iAs exposure alters cardiomyocyte Ca^2+^ handling and shortening

We observed an increase in heart size without a change in *in vivo* function as measured by echocardiography, so we next examined the function of isolated ventricular cardiomyocytes. Ca^2+^ transients and sarcomere shortening were assessed in cardiomyocytes isolated at P14 and P15 from postpartum dam hearts exposed to iAs, along with non-exposed postpartum controls, non-pregnant non-exposed controls and non-pregnant mice exposed to iAs. We found that gestational exposure to 100 µg/L iAs significantly increased the time to peak of the Ca^2+^ transient for both postpartum and non-pregnant female mice (Fig. 3A). Ca^2+^ transient amplitude was significantly decreased in cardiomyocytes from both exposed and non-exposed postpartum females vs. their non-pregnant controls, but cardiomyocytes from iAs exposed postpartum female mice showed a further significant reduction in Ca^2+^ transient amplitude compared to non-exposed postpartum female mice (Fig. 3B). Cardiomyocytes from postpartum females exposed to 100 µg/L iAs also showed a significant increase in Ca^2+^ transient relaxation time to baseline at 50% (Fig. 3C), as well as increased relaxation tau (Fig. 3D) compared to non-exposed postpartum females and non-pregnant iAs exposed mice.

**Figure 3:**
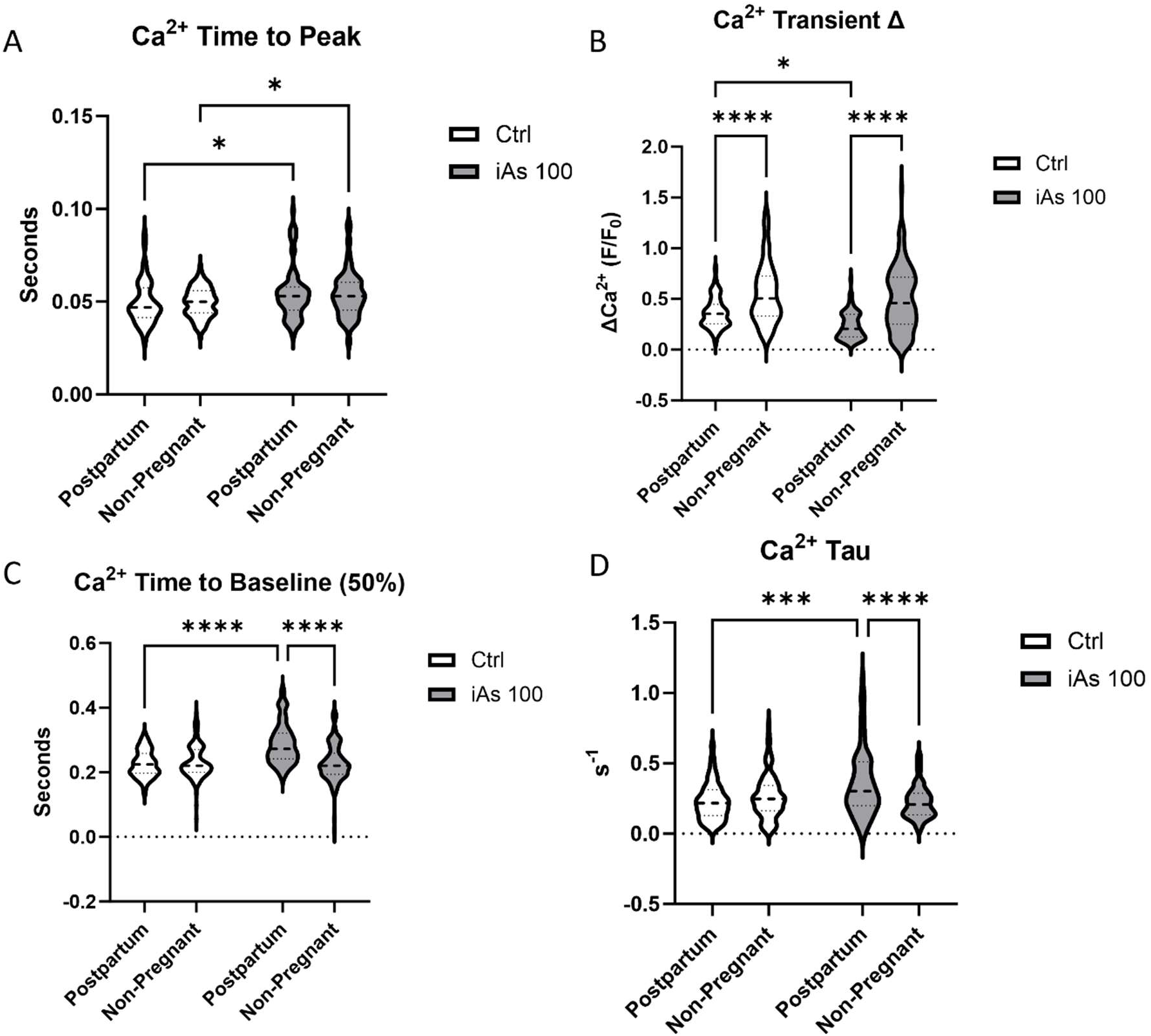
Gestational iAs exposure alters Ca^2+^ handling in the cardiomyocyte. Time to peak (A), transient amplitude (B), time to baseline (C) and tau (D) for Ca^2+^ transients as measured in individual cardiomyocytes from iAs-exposed or non-exposed postpartum female hearts (*n*= 40-80 myocytes/heart from 4 hearts/group); *p<0.05, **p<0.01, ***p<0.001, ****p<0.0001 vs. control). Outliers were determined by the ROUT method (Q=1%) and significance was determined by 2-way ANOVA.

Additionally, we found a significant increase in the time to peak sarcomere shortening in cardiomyocytes from postpartum females exposed to 100 µg/L iAs compared to non-exposed postpartum controls and 100 µg/L iAs exposed non-pregnant female mice (Fig. 4A). We also found that while percent sarcomere shortening was significantly increased in cardiomyocytes from non-exposed postpartum female mice vs. non-exposed, non-pregnant female mice, this was not the case in cardiomyocytes from postpartum females exposed to 100 µg/L iAs; we observed a significant decrease in percent shortening compared to non-exposed postpartum controls, and no difference when comparing non-pregnant and postpartum female mice exposed to 100 µg/L iAs (Fig. 4B). Percent sarcomere shortening was significantly increased with exposure to 100 µg/L iAs in non-pregnant mice compared to non-exposed, non-pregnant mice. Cardiomyocytes from postpartum females exposed to 100 µg/L iAs also showed a significant increase in sarcomere time to baseline at 50% compared to non-exposed postpartum controls, and non-pregnant mice exposed to 100 µg/L iAs (Fig. 4C). There was an opposite effect on tau when considering postpartum and non-pregnant comparisons within exposure groups. Sarcomere tau was significantly increased in 100 µg/L iAs exposed postpartum female mice compared to iAs exposed non-pregnant and non-exposed postpartum controls. However, there was a significant decrease and an opposite effect on tau when comparing non-exposed postpartum mice and non-pregnant 100 µg/L iAs exposed female mice to non-pregnant, non-exposed female mice (Fig. 4D).

**Figure 4:**
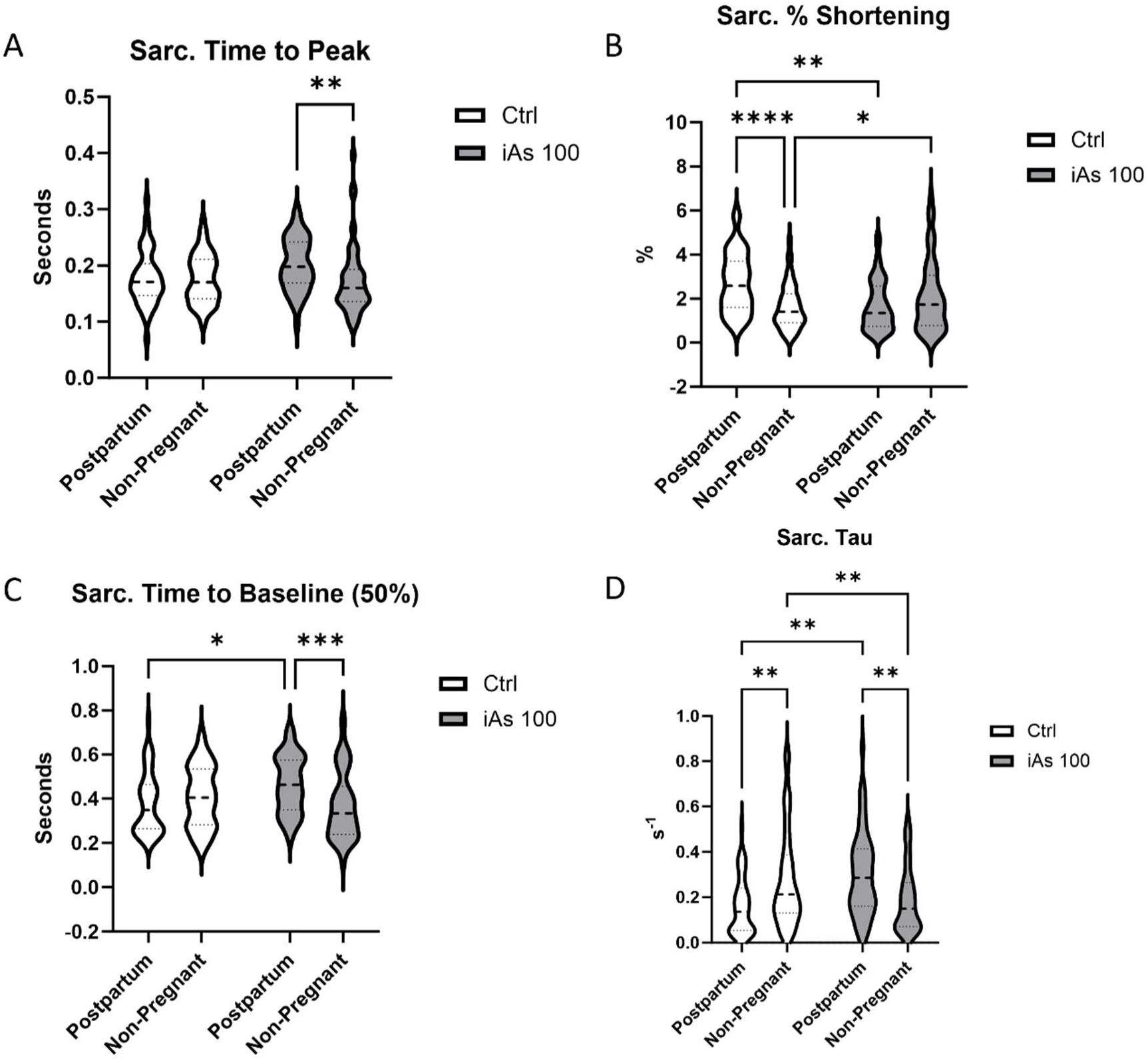
Gestational iAs exposure alters sarcomere shortening in the cardiomyocyte. Time to peak (A), percent shortening (B), time to baseline (C), and tau (D) for sarcomere shortening as measured in individual cardiomyocytes from iAs-exposed or non-exposed postpartum female hearts (*n*= 40-80 myocytes/heart from 4 hearts/group); *p<0.05, **p<0.01, ***p<0.001, ****p<0.0001 vs. control). Outliers were determined by the ROUT method (Q=1%) and significance was determined by 2-way ANOVA.

### iAs exposure and Ca^2+^ handling / contractile protein expression

To determine if the observed changes in cardiomyocyte function were a result of transcript or protein expression differences, we next probed the expression of critical regulators of Ca^2+^ handling and cardiomyocyte contractile proteins. We found that gestational exposure to 100 µg/L iAs did not alter transcript levels for the sarco/endoplasmic reticulum ATPase 2a (SERCA2a, *Atp2a2*), the α subunit of the L-type Ca^2+^ channel (LTCC, *Cacna1d*) or the ryanodine receptor (RyR, *RyR2*) (Fig. 5A), or the protein expression of SERCA2a or the α-2 subunit of the LTCC in postpartum female mice (Fig. 5B). We also found no change in total phospholamban (PLBN) expression (Fig. 5B) or phosphorylated phospholamban (p-PLBN) with exposure to 100 µg/L iAs (Fig. 5B). However, we did find a significant decrease in the protein expression of cardiac myosin binding protein C (cMyBP-c), as well as a significant reduction in α-actin, in postpartum hearts after 100 µg/L iAs exposure (Fig. 5C).

**Figure 5:**
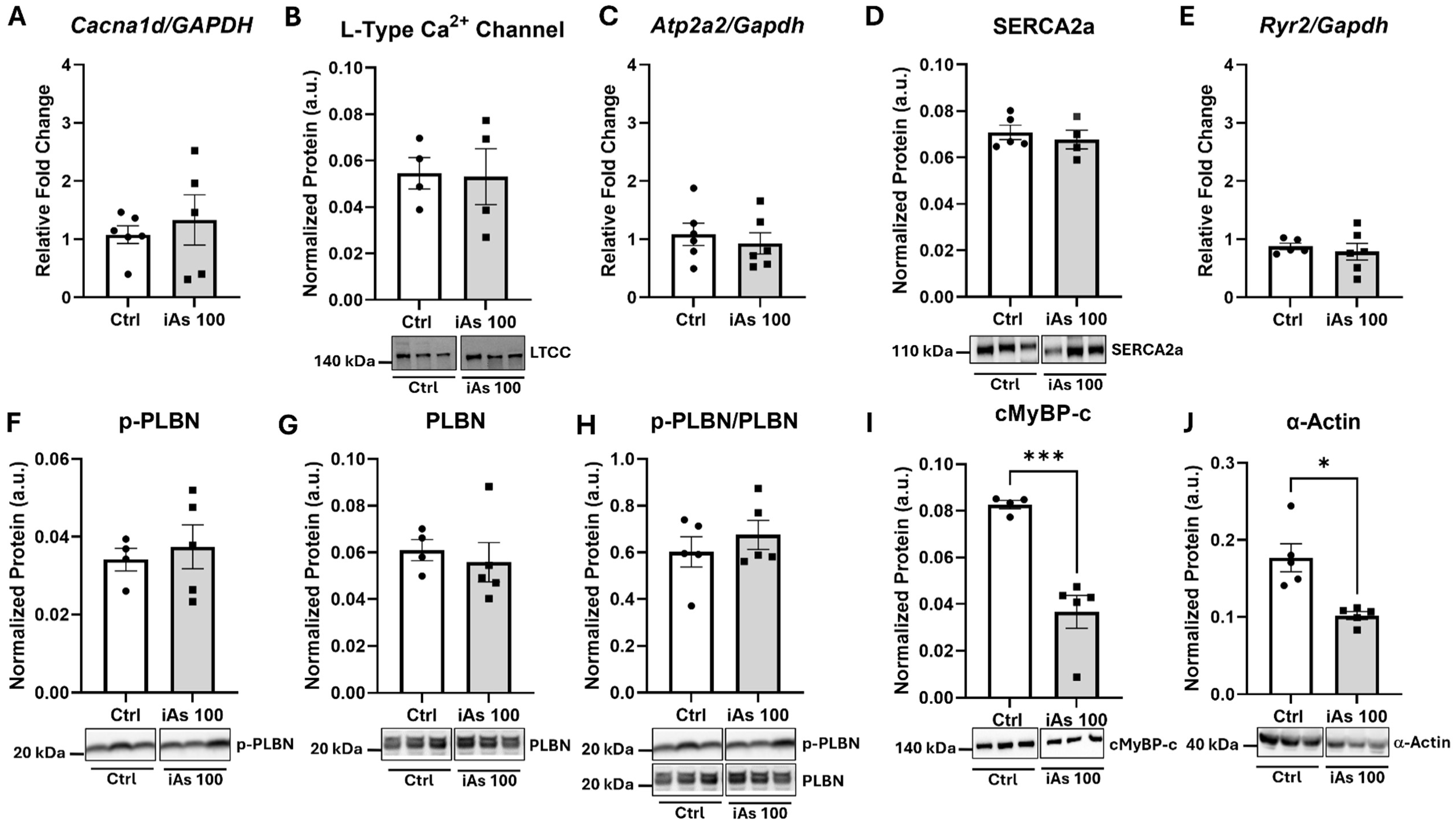
Gestational iAs exposure alters expression of contractile proteins at P12. Myocardial expression of the L-type Ca^2+^ channel (*Cacna1d*, LTCC) (A-B), the sarco/endoplasmic reticulum Ca^2+^ ATP-ase (*Atp2a2*, SERCA2a) (C-D), the ryanodine receptor (*RyR2*) (E), phosphorylated phospholamban (p-PLBN) (F), total phospholamban (PLBN) (G), phosphorylated/total phospholamban (H), cardiac myosin binding protein C (cMyBP-c) (I), and α-actin (J) in iAs-exposed or non-exposed postpartum females. (*n*=4-6 hearts/group; *p<0.05, ***P<0.001 vs. control). Significance was determined by Mann-Whitney.

## DISCUSSION

We report here for the first time that iAs exposure during gestation increases postpartum heart size, induces Ca^2+^ handling dysregulation and contractile changes at the level of the cardiomyocyte, and alters the expression of key contractile proteins. We found that gestational exposure to 100 or 1000 µg/L iAs increased postpartum heart size at P12, but cardiac function was not altered at P5 (Fig. 1). We next examined markers of hypertrophy and found that gestational iAs exposure significantly downregulated markers of physiologic hypertrophy, including Akt and p-eNOS (Fig. 2). We also found changes in Ca^2+^ handling (Fig. 3) and shortening (Fig. 4) at the level of the isolated cardiomyocyte, including decreased Ca^2+^ transient amplitude and increased relaxation time. Correspondingly, we observed a reduction in sarcomere shortening and increased sarcomere relaxation time and tau. To better understand these changes in cardiomyocyte function, we also examined the expression of select proteins and found a decrease in both cMyBP-c and α-actin expression (Fig. 5).

Our prior study showed that iAs blunted maternal cardiac growth during pregnancy, while our current study examines potential impacts on the postpartum maternal mouse heart^35^. Importantly, our work demonstrates that gestational iAs exposure induces a mechanistic dysregulation in Ca^2+^ handling and contraction at the level of the isolated cardiomyocyte that extends beyond pregnancy and the iAs exposure window. Furthermore, our isolated cardiomyocyte findings result from the combination of pregnancy and 100 µg/L iAs exposure, indicating that in females, iAs exposure alone is not sufficient to induce this dysfunctional phenotype. This is consistent with our prior studies, which showed alterations to cardiac structure and function with long-term exposure to iAs or cadmium in male mice, but no apparent pathology in non-pregnant females^21,22,42^. However, in the current study, we found pregnant females to be especially sensitive to cardiac alterations after gestational iAs exposure, and as such, our findings suggest that the combined stress of pregnancy and iAs exposure leads to altered heart growth and signaling which persist after pregnancy and the exposure period. Importantly, the effect of iAs exposure on the postpartum maternal heart has not been examined in human pregnancy, and furthermore, studies on the molecular mechanisms that underpin maternal cardiac hypertrophy during pregnancy and its postpartum reversal are severely lacking. As such, our study is the first to elucidate the potential effects of an environmentally relevant iAs exposure on the maternal cardiovascular system after pregnancy, and our findings underscore the importance of additional studies in humans to uncover potential cardiotoxic effects.

### Gestational iAs exposure increases heart size postpartum

Cardiac hypertrophy is an essential process which occurs during pregnancy. Importantly, under normal physiologic conditions, the heart should remodel back to its original size postpartum^23,26^. Cardiac hypertrophy is generally classified as physiologic or pathologic, with a primary distinction being the reversibility of physiologic hypertrophy ^43^. Physiologic cardiac hypertrophy occurs during pregnancy due to plasma volume expansion and hormonal changes^44,45^, which induce pressure overload in the heart. In mice, pregnancy-induced cardiac hypertrophy can reverse as early as P7^26^, but we found that in females exposed to iAs postpartum heart size was increased at P12 (Fig. 1a), potentially indicating that iAs exposure during gestation alone is sufficient to alter signaling pathways postpartum. Interestingly, in a prior study, we found that iAs exposure during pregnancy reduced cardiac growth relative to controls at E17.5^35^. However, rather than reversing cardiac growth postpartum, our current findings suggest that gestational iAs exposure triggers the enlargement of the maternal heart postpartum or prevents effective reverse remodeling. Furthermore, while mechanisms underlying the postpartum reversal of pregnancy-induced cardiac growth are not well studied, hormonal signaling and reduced preload are thought to play an important role^46^. Litter size has been shown to alter maternal blood pressure in rodents, but our prior study identified no changes in litter size after gestational iAs exposure, suggesting that litter size is unlikely to contribute to our phenotype^46,47^. Additionally, we previously identified reduced estrogen receptor α expression in the heart at E17.5 with gestational iAs exposure, supported a potential role for altered hormonal signaling in our observed increase in postpartum heart size after gestational iAs exposure^35^. Therefore, future studies will investigate preload and hormonal signaling in the postpartum maternal heart following gestational iAs exposure.

To understand how increased heart size affects cardiac function and structure, transthoracic echocardiography was performed on postpartum dams. Physiologic hypertrophy, observed during pregnancy, would traditionally increase functional parameters, such as cardiac output and stroke volume^24,48^, but in the current study, we did not observe any changes in these cardiac parameters (Fig. 1). This suggests that the increase in cardiac size may not be beneficial in terms of heart function. An increase in postpartum heart size in tandem with preserved cardiac function may indicate that gestational iAs exposure induces a form of cardiac remodeling at P5 that may either escalate into cardiac pathology as the postpartum period progresses or predispose the postpartum heart to cardiac dysfunction under stress. Thus, it will be important for future studies to examine cardiac function at later postpartum timepoints to understand the long-term consequences of gestational iAs exposure on maternal cardiac function.

### iAs decreases p-eNOS and Akt postpartum

To understand the observed increase in heart size, we next probed proteins associated with physiologic or pathologic hypertrophic signaling. Physiologic and pathologic hypertrophy have distinct signaling pathways with some overlap^43,49^. Pathologic hypertrophy is not reversible and ultimately culminates in heart failure, and is associated with a switch in myosin isoforms from α-myosin (*Myh6*) to β-myosin (*Myh7*)^50,51^, which we did not observe. Physiologic hypertrophy, on the other hand, is reversible and has major drivers such as protein kinase B (Akt)^52^. ERK1/2 specifically plays an important role in cardiac hypertrophy but has been found to be upregulated with certain forms of both physiologic and pathologic hypertrophy^53–55^. Specifically, in pregnancy ERK1/2 is thought to be beneficial, as the ERK pathway is activated by both estrogen and the onset of hypertrophy with pressure overload^56^. We probed for proteins traditionally associated with physiologic and pathologic hypertrophy and found that iAs exposure decreased eNOS activation at its activating phosphorylation site (S1177) and decreased overall Akt expression (Figs. 3B and 3C). Akt is traditionally upregulated in physiologic hypertrophy and activates growth pathways, while eNOS upregulates angiogenesis which needed for increased cardiac growth^57,58^. While we expected to see a clear increase in markers for pathologic or physiologic hypertrophy, we instead observed a decrease in markers known to be strongly associated with physiologic hypertrophy, and more specifically pregnancy-induced hypertrophy after gestational exposure to 100 µg/L iAs.

We found that heart size was increased in postpartum hearts exposed to 100 µg/L iAs during gestation, but it is important to understand if this hypertrophy may be characterized as pathologic or physiologic. In the case of our current findings, we may be observing an intermediate phenotype which does not clearly fit into either category. The downregulation of traditional physiologic markers of hypertrophy may indicate that our observed postpartum cardiac hypertrophy is not physiologic. However, this characterization is neither exhaustive nor conclusionary, and more mechanistic experimentation is needed to fully understand the signaling behind the phenotype. Our timepoints may also be too early in the postpartum period to fully categorize these changes, so longer-term studies on both function and mechanism are needed.

### iAs exposure alters cardiomyocyte Ca^2+^ handling

We next wanted to understand the impact of gestational iAs exposure on cardiomyocyte function by examining Ca^2+^ transients and sarcomere shortening postpartum. Interestingly, we found that at P14/15, gestational exposure to 100 µg/L iAs significantly altered Ca^2+^ transients in postpartum females compared to non-exposed postpartum controls and non-pregnant females previously exposed to iAs. While we found no changes in cardiac function as measured by *in vivo* echocardiography, these measurements were done at P5, so it is possible that these changes were not yet apparent. When comparing both postpartum control and non-exposed, non-pregnant female mice to iAs exposed postpartum females and exposed non-pregnant females which had been exposed to iAs for the same time frame, we found that the time to peak for the Ca^2+^ transient was equally affected by pregnancy regardless of iAs exposure. To our knowledge, this is the first study to report that Ca^2+^ transient time to peak increases at P14/15 in the mouse, potentially indicating slower release of Ca^2+^ after the action potential. We also found that irrespective of exposure, postpartum female mice had significantly smaller Ca^2+^ transient amplitude compared to non-pregnant female mice, indicating that there is less Ca^2+^ released into the cytosol during contraction at P14/15. During pregnancy there are increased calcium transients in the heart through an ERα induced mechanism^59^. While our study did not examine transients during pregnancy, our findings indicate that calcium transients during pregnancy may sharply decrease after pregnancy resulting in levels that are lower than non-pregnant females. Together, we found that iAs exposure during pregnancy reduced Ca^2+^ transient amplitude and prolonged the relaxation of the Ca^2+^ transient. Furthermore, the combination of iAs exposure and pregnancy induced alterations to Ca^2+^ handling at the level of cardiomyocyte that persisted beyond pregnancy and the iAs exposure window. These changes were also not due to pregnancy or iAs exposure alone, indicating that pregnancy represents a critical period for iAs exposure.

Consistent with our changes in cardiomyocyte Ca^2+^ handling, we also found that exposure to 100 µg/L iAs resulted in alterations to sarcomere shortening in isolated cardiomyocytes. We found that female mice exposed to iAs during gestation had increased time to contraction after stimulation. We also found in iAs exposed postpartum females compared to non-exposed postpartum controls that the isolated cardiomyocyte is re-lengthening more slowly, despite the decrease in overall contraction. These results are congruent with our observed alterations to Ca^2+^ handling in iAs-exposed animals, and we would expect that observed changes to sarcomere contraction are due to alterations in Ca^2+^ handling. Interestingly, while we see changes at the level of the individual cardiomyocyte at P14/15, we observed no change in whole heart function at 5, indicating that the changes we observe at P14/15 may occur after the P5 timepoint. At the level of the whole heart, we would expect a reduction in ejection fraction or fractional shortening. However, there are compensatory mechanisms within the whole heart which may counteract our observed cellular changes. This discrepancy between whole heart and isolated cardiomyocyte function may also be due to the lack of a mechanical load in isolated cardiomyocytes. A mechanical load can stretch the cardiomyocyte, referred to as preload, thus allowing for the cardiomyocyte to generate more force^60^. As such, this lack of preload may affect how isolated cardiomyocytes behave and contract in isolation^60^ and therefore, in our experiments, the observed changes in sarcomere shortening could be a direct result of alterations to Ca^2+^-handling. Changes to Ca^2+^ sensitivity may also explain this discrepancy, as iAs exposure has been shown to affect intracellular Ca^2+61,62^. Epidemiologic studies have also reported an association between iAs exposure and hypertension^18,63^, which is thought to occur with an iAs-induced increase in Ca^2+^ sensitization and hypercontraction in blood vessels^64^. Since we observed less Ca^2+^ release into the cytosol in the isolated cardiomyocyte, it may indicate that less Ca^2+^ is needed to induce contraction, providing further evidence that iAs exposure induces Ca^2+^ sensitization. However, additional studies are needed to confirm this mechanism.

### iAs exposure and excitation-contraction protein expression

Since we observed changes in cardiomyocyte Ca^2+^ handling and contraction, we next probed for proteins important for excitation-contraction coupling to determine potential iAs-induced impacts. Contractility may be affected by either a change in Ca^2+^ handling or myofilament proteins. We probed proteins associated with Ca^2+^ handling, including SERCA2a, phospholamban and the LTCC, but found no changes in the expression of these proteins (Fig. 5). We also found no difference in transcript levels for RyR (Fig. 5), which is responsible for the release of Ca^2+^ from the sarcoplasmic reticulum during excitation-contraction coupling. However, since RyR activity can be regulated in numerous ways, including via reactive oxygen species^65,66^ which are known to be generated with iAs exposure, future studies will also need to examine the activity of RyR, and other Ca^2+^-handling proteins like SERCA2a and LTCC.

Furthermore, we characterized the expression of the myofilament proteins cMyPB-c and α-actin, both of which showed decreased expression with gestational iAs exposure (Fig. 5). cMyPB-c is a protein associated with myosin that is localized to the crossbridge-containing zone of the sarcomere,^67^ playing a crucial role in contraction of the cardiomyocyte. cMyBP-c binds myosin to regulate contraction, and reduces actomyosin ATPase activity until it is phosphorylated, at which point it releases the “brake” on crossbridge cycling and allows for contraction^67^. Phosphorylation can occur in response to β-adrenergic stimulation via PKA or CaMKII^68–72^. When cMyBP-c is partially extracted or removed in a homozygous knockout mouse, there is an increase in Ca^2+^ sensitivity, force of contraction, and time to relaxation in the cardiomyocyte^73–76^. It is possible that the reduction in cMyBP-c may influence contraction speed independent of Ca^2+^. As mentioned previously, it is also possible that iAs may alter Ca^2+^ sensitivity, as we do see an increase in time to relaxation in the cardiomyocyte. Therefore, it is possible that gestational iAs exposure has the capacity to decrease cMyPB-c protein expression after pregnancy, thereby altering Ca^2+^ sensitivity and cardiomyocyte contraction. α-Actin is the thin filament protein in the contractile unit which allows for myosin binding and contraction. Reduction in both α-Actin and cMybp-c have been implicated in cardiomyopathy and cardiac disease^77^. However, future experiments will focus on the exact mechanism by which this occurs, as well as future endpoints to see if these changes persist.

## LIMITATIONS

While this study provides important insights into the effects of iAs on maternal health postpartum, there are limitations that require acknowledgement. Firstly, this study uses limited postpartum timepoints, so future studies should examine multiple extended timepoints to understand how gestational iAs exposure affects maternal cardiac health long-term. Additionally, echocardiography was only performed at one timepoint and under anesthesia, and thus does not provide a complete picture of the changing postpartum cardiac landscape. We also recognize that echocardiography was performed prior to terminal endpoints, and there are time-dependent effects on cardiac function which may not have been captured. Additionally, we performed standard M-mode echocardiography, as opposed to tissue Doppler or two-dimensional speckle tracking to assess, so we were not able to capture measures such as strain. We also did not perform additional techniques to fully evaluate heart physiology, including pressure-volume loops or telemetry, but these techniques will be employed in future studies. Furthermore, exposure was started at E2.5 during gestation, while a real-world exposure would encompass a lifetime. Although this is a limitation of our study, this exposure scenario is important for defining a critical window of maternal exposure. Experiments were also conducted using a mouse model, and while mice are well characterized and commonly utilized for studies examining pregnancy and iAs exposure, it is important to note that iAs metabolism in a mouse is different in mice compared to humans^6,78,79^. The mouse metabolizes iAs more efficiently than humans, making higher doses in mice equivalent to lower doses in human populations. There are also key differences in murine pregnancy compared to that of humans. Finally, we only examined two iAs doses in our study, 100 µg/L and 1000 µg/L. Lower doses should be used in future studies to understand if these effects persist at lower doses, such as the recommended limit of 10 µg/L provided by the Environmental Protection Agency.

## CONCLUSION

iAs exposure during gestation alone resulted in an increase in postpartum heart size and altered Ca^2+^-handling and contraction at the level of the cardiomyocyte. Importantly, impacts to heart size and cardiomyocyte function were noted after pregnancy and the iAs exposure window, suggesting that the impact of gestational iAs exposure is persistent. Furthermore, at the molecular level, we found that the expression of multiple proteins associated with excitation-contraction coupling to be altered with gestational iAs exposure. Taken together, our results underscore the potential for gestational iAs exposure to have lasting impacts on the maternal heart beyond the iAs exposure period and since this is an understudied area in human pregnancy, it is critical that future human exposure studies examine the maternal heart during and after pregnancy. Furthermore, this research adds to a growing body of evidence implicating iAs as a driver of cardiovascular disease, and further suggests that the environment plays a role in the etiology of cardiovascular-related adverse pregnancy events. These findings further emphasize the importance of effective drinking water regulation.

## ACKNOWLEDGEMENTS

We acknowledge the technical assistance of the Small Animal Cardiovascular Phenotyping and Model Core at the Johns Hopkins University School of Medicine.

## AUTHOR CONTRIBUTIONS

N. Taube; conceived and designed research, performed experiments, analyzed data, interpreted results of experiments, prepared figures, drafted manuscript, edited and revised manuscript, approved final version of manuscript. M. Steiner; performed experiments, analyzed data, interpreted results of experiments, edited and revised manuscript, approved final version of manuscript. O. V. Ebenebe; designed research, performed experiments, analyzed data, interpreted results of experiments, edited and revised manuscript, approved final version of manuscript. R. Kabir; performed experiments, edited and revised manuscript, approved final version of manuscript. H. Garbus; performed experiments, edited and revised manuscript, approved final version of manuscript. S.M. Alam El Din; performed experiments, edited and revised manuscript, approved final version of manuscript. E. Illingworth; performed experiments, edited and revised manuscript, approved final version of manuscript. N. Wang; performed experiments, edited and revised manuscript, approved final version of manuscript. B.L. Lin; performed experiments, analyzed data, interpreted results of experiments, edited and revised manuscript, approved final version of manuscript. M. Kohr; conceived and designed experiments, interpreted results of experiments, provided support and funding, edited and revised manuscript, approved final version of manuscript.

